# Lesion Localization of Time Disorientation in Patients With Focal Brain Damage

**DOI:** 10.1101/2022.05.24.493338

**Authors:** J. Skye, J. Bruss, G. Herbet, D. Tranel, AD. Boes

## Abstract

**Background and Objectives:** Time orientation is a fundamental cognitive process in which one’s personal sense of time is matched with a universal reference. Assessment of time orientation is a ubiquitous component of neurological mental status examinations and neuropsychological assessments, yet its neural correlates remain unclear. Large bilateral lesions have been associated with deficits in time orientation, but more specific regions of the brain implicated in time disorientation following focal unilateral damage are relatively unknown. The current study investigates the anatomy of time disorientation and its network correlates in patients with focal brain lesions.

**Methods:** 550 patients with acquired, focal brain lesions participated in this study, identified retrospectively from the Iowa Neurological Patient Registry. Time orientation was assessed 3 months or more after lesion onset using the Benton Temporal Orientation Test (BTOT), and 39 patients were identified as having chronic impairment in time orientation defined as a score of 3 or worse on the BTOT. Multivariate lesion-symptom mapping and lesion network mapping were used to evaluate the anatomy and networks associated with time disorientation. Performance on a variety of neuropsychological tests was compared between the time oriented and time disoriented group.

**Results:** 39 patients were identified as having chronic impairment in time orientation. Multivariate lesion-symptom mapping showed that lesions of the posterior cortices were associated with impaired time orientation, including medial temporal lobes, occipitotemporal cortex, and precuneus (r=0.21, p<.001). Individuals with time disorientation tended to have concomitant impairments in memory, visuospatial ability, and naming. Follow-up analyses of individuals with unilateral lesions and those with relatively unimpaired cognition in other domains implicated the precuneus and parahippocampal gyrus in time orientation. Lesion network mapping demonstrated that these regional findings occurred at nodes of the default mode and visual networks. Individuals with time disorientation tended to have concomitant impairments in memory, visuospatial ability, and naming.

**Discussion:** We interpret these findings as novel evidence for the role of posteromedial cortices extending from the precuneus to the medial temporal lobe in supporting time orientation.

## Introduction

Humans perceive time as flowing continuously from past through present to future. Time orientation describes the ability to match one’s internal representation of time to a universal reference system, or ‘clock time,’ on the order of hours to years [1, 2]. Assessing time orientation is a fundamental component of mental status exams, usually tested along with orientation to person and place. There are valid, standardized methods of assessing time orientation in clinical settings such as the Benton Time Orientation Task (BTOT) which asks five questions about the time of the day, day of the week, and date (day of month, month, and year) [1]. Time disorientation can occur in a wide variety of psychiatric and neurological disorders [3]. Identifying this impairment, sometimes referred to as chronotaraxis, is useful in understanding a patient’s overall cognitive status. Moreover, time disorientation is associated with other aspects of cognition including planning [4], decision making [5], and memory [6, 7]. Time orientation is frequently assessed, yet very little is known about the neural substrates that support this function. This study investigates time disorientation following focal, acquired brain lesions.

The anatomical systems most critical to representing the passage of time and associating it with clock time are not well understood. Elucidating the anatomy is particularly challenging because, unlike many basic perceptual systems, there is no clear sensory receptor for time: the source of information the brain uses to estimate time is unclear, but presumed to be multimodal. There is a robust association of time disorientation and amnesia, which highlights a potential role for medial temporal lobe (MTL) structures such as the hippocampus and adjacent cortices [7, 8]. “Time cells” sensitive to duration have been identified in the MTL in rodents and humans [8-10], and degeneration of this system is associated with worsening time disorientation in Alzheimer’s disease. Yet time disorientation cannot be explained by amnesia alone [11], with some studies suggesting that the temporality and content of memory are distinct [12, 13]. Other regional anatomy that has been implicated in time orientation includes the mediodorsal nucleus of the thalamus [14, 15], the right cerebral hemisphere [16], the precuneus [17], and the prefrontal cortex [17]. Whether specific brain networks support time orientation is relatively unknown, though some prior work has identified a potential role for the default mode network, which aligns with the above-cited studies of MTL and precuneus involvement. The extent to which these regional associations are correlational or causal is unclear, as is the relative contribution of each region.

Studying patients with focal acquired brain lesions who are impaired in time orientation has the potential to aid in the elucidation of brain regions most critical for supporting time orientation. To date most lesion studies of time disorientation have been limited by small sample sizes, focusing on a specific brain region (e.g., the thalamus), or including multifocal lesions from progressive disorders or traumatic brain injury [14, 15, 18, 19]. Another challenge to this line of research is time disorientation after brain injury is relatively common in the acute epoch, but spontaneously resolves in ∼95% of patients within about six weeks [20]. However, in a select group of individuals, time disorientation persists to become a chronic impairment lasting 3 or more months after lesion onset, presumably due to lesions of anatomical structures and networks that are critical in supporting time orientation. To the best of our knowledge, a large-scale lesion-symptom mapping study of time disorientation using modern lesion mapping methods has not been performed. In addition to investigating lesion location, lesions and associated symptoms can also be interpreted in the context of disrupting a broader network that extends beyond the anatomical boundaries of the lesion [21]. Lesion network mapping is an approach that combines traditional lesion-symptom mapping approaches with functional connectivity information derived from large normative ‘connectome’ datasets to evaluate lesion associated networks. This approach has been useful for uncovering network correlates of a number of lesion syndromes that have been historically challenging to ‘localize’ [21-24] but has not been applied to time orientation to date.

Here, we utilize a large registry of 550 patients with focal, acquired brain lesions, who have standardized time orientation assessments performed more than 3 months after the lesion onset. Our goals were to: 1) perform a data-driven analysis evaluating the regional anatomy associated with time disorientation, 2) perform lesion network mapping to evaluate whether there is a network organization of any of the findings in #1, and 3) evaluate the cognitive profile of individuals with time disorientation in order to determine whether other cognitive impairments (such as memory deficits) co-occur systematically with time disorientation.

## Methods

We used neuropsychological test data and neuroimaging data available through the Iowa Neurological Patient Registry to perform our analyses. Patients in the Registry gave informed, written consent prior to participation in this study, and the study was approved by the Institutional Review Board of the University of Iowa. Registry patients underwent comprehensive neuropsychological evaluation in accordance with Benton Neuropsychology Laboratory protocols [25], which included an extensive battery of neuropsychological tests designed to assess major domains of cognition and behavior. Data were collected during the chronic epoch of recovery (>3 months after lesion onset) in patients over the age of 17. Prior to enrollment in the Registry, patients are screened for pre-existing psychiatric and neurological conditions. For the current analysis, in order to ensure that measured time disorientation was not confounded by receptive language deficits, patients were excluded from this study if they had impaired scores on the Boston Diagnostic Aphasia Examination (BDAE) Complex Ideational Material Test, the Multilingual Aphasia Examination (MAE) Token Test, or the MAE Aural Comprehension Test (defined as raw scores ≤ 8, ≤ 36, or ≤ 14, respectively). Patients with focal, acquired brain lesions were included regardless of etiology.

We used the BTOT to measure time orientation [26, 27]. In this test, the examiner verbally asks the patient five questions about where they are in time (*Table 1*). A total of 625 patients were given a score on a scale from 0 to -113 with 0 being considered unimpaired performance. The BTOT score obtained most contemporaneously with the date of the patient’s research brain scan was used if a patient had been administered the BTOT more than once. Patients with a score of 0 constitute the brain-damaged comparison group (n=511). Following the normative standards for the BTOT [26], patients in our study with a score of -3 or worse were considered impaired; there were 39 such patients (mean score = -15.10, standard deviation = -22.33, range = -3 to -102). We excluded 75 patients with a score of -1 or -2, which created a greater contrast between the impaired and unimpaired groups; these scores correspond to common errors healthy adults make in self-referential timing (e.g., being 30 minutes off for the current time or being a day off for the day of the week), and are not “impaired” according to the normative standards for the BTOT. 550 participants met the criteria for this study (*Table 2*). We used the Wilcoxon Rank Sum test to compare the age, education, and lesion volume between time oriented and time disoriented patients.

**Table 1.** The Benton Temporal Orientation Test (BTOT) asks five questions about where a person is in time. If an incorrect response is provided, a specified number of points is deducted from a starting score of zero, according to the nature of their error. Each time interval has a maximum deduction for that item.

| **Time Interval** | **Scoring Criteria** | **Maximum Error** |
| --- | --- | --- |
| month | -5 points per month off | -30 points |
| day of the month | -1 points per day off | -15 points |
| year | -10 points per year off | -60 points |
| day of the week | -1 points per day off | -3 points |
| time of day | -1 points per half hour off | -5 points |

**Table 2.** Demographics information for patients in the study.

| Demographics |  |  |  |  |
| --- | --- | --- | --- | --- |
| Characteristic |  | All Patients<br>(N=550) | Impaired<br>(n=39) | Unimpaired<br>(n=511) |
| Age in years | mean (std) | 52.42 (14.46) | 59.71 (14.83) | 51.84 (14.30) |
| Education in years | mean (std) | 13.40 (2.75) | 13.18 (3.15) | 13.42 (2.72) |
| Gender | female | 268 | 19 | 249 |
|  | male | 282 | 20 | 262 |
| Handedness | right | 493 | 34 | 459 |
|  | left | 41 | 4 | 37 |
|  | mixed | 16 | 1 | 15 |
| Race | African American | 4 | 0 | 4 |
|  | American Indian | 3 | 0 | 3 |
|  | Caucasian | 541 | 39 | 502 |
|  | other/unknown | 2 | 0 | 2 |
| Ethnicity | Hispanic | 2 | 0 | 2 |
|  | non-Hispanic | 547 | 39 | 508 |
|  | unknown | 1 | 0 | 1 |
| Lesion volume in mm<sup>3</sup> | mean (std) | 37774.76<br>(56451.84) | 64523.69<br>(76716.88) | 35710.29<br>(54208.70) |
| Lesion laterality | right | 205 | 13 | 192 |
|  | left | 228 | 7 | 221 |
|  | bilateral | 117 | 19 | 98 |
| Etiology | ischemic stroke | 287 | 17 | 270 |
|  | hemorrhage | 119 | 7 | 112 |
|  | benign tumor resection | 78 | 4 | 74 |
|  | other resection | 29 | 3 | 26 |
|  | focal contusion | 17 | 3 | 14 |
|  | herpes simplex or limbic encephalitis | 12 | 5 | 7 |
|  | multiple etiologies | 8 | 0 | 8 |

Structural neuroimaging was obtained 3 months or more after the lesion onset and the boundaries of the lesions were manually segmented for all 550 scans using standard procedures [28]. Lesion masks for neuroimaging data acquired prior to 2006 were generated using the MAP-3 method [28, 29] wherein the boundaries of the lesion are traced onto a template brain. Lesions in neuroimaging data acquired in 2006 and after were manually traced onto the patient’s T1 native scan in FSL [30] and then subsequently transformed into MNI152 space. The anatomical accuracy of the native trace and the transformed lesion mask were confirmed and edited if necessary by a neurologist (A.D.B.) who was blind to all demographic and cognitive data.

Lesion-symptom mapping was used to identify regions of damage associated with time disorientation. LESYMAP, a multivariate lesion-symptom mapping technique that uses sparse canonical correlation analysis for neuroimaging (SCCAN) was employed using BTOT data [31]. We first performed lesion-symptom mapping of all 550 subjects using BTOT score as a binary variable. We also performed an analysis that was limited to subjects with unilateral lesions. Prior studies have reported large bilateral lesions from etiologies such as encephalitis are associated with time disorientation [32], but it can be difficult to interpret which regional findings are most critical when lesions are large and bilateral. We performed lesion-symptom mapping after removing all subjects with bilateral lesions in an effort to evaluate whether time disorientation also occurs from unilateral lesions, and if so, which regions are implicated. The lesion-symptom mapping analysis of unilateral lesions was performed by representing lesions according to their true laterality and also by flipping half the lesions on the x-axis to a single hemisphere. This latter analysis maximizes statistical power, but is limited in that it assumes no laterality differences and is unable to detect them if they do exist.

Lesion network mapping was performed similarly to prior work [22, 33], in which networks were derived from brain areas representing the strongest brain-behavior associations identified from the lesion-symptom map. This differs from the approach of using each lesion mask to “seed” the network analysis [21] to avoid some of the problems associated with signal averaging from large lesions [34]. A 4mm spherical region of interest (ROI) was placed at regional peaks with the highest voxel intensity, representing the strongest regional association with time disorientation as identified by the lesion-symptom mapping analysis and used to seed lesion network maps. Resting state functional connectivity MRI (rs-fcMRI) data from a normative database was used, as in previous work (n=98 healthy right-handed subjects; 48 male subjects, age 22 +/- 3.2 years) [35]. The rs-fcMRI data were processed in accordance with previously described methods [36]. Briefly, participants completed two 6.2-minute resting state functional MRI scans during which they were asked to rest in the scanner (3T, Siemens) with their eyes open (TR = 3,000 ms, TE = 30 ms, FA = 85°, 3 mm voxel size [27 mm^3^], FOV = 216, 47 axial slices with interleaved acquisition and no gap). Functional data were acquired at 3 mm voxel size and spatially smoothed using a Gaussian kernel of 4 mm full-width at half-maximum. The data were temporally filtered (0.009 Hz < f < 0.08 Hz) and several nuisance variables were removed by regression, including (a) six movement parameters computed by rigid body translation and rotation during preprocessing, (b) mean whole brain signal, (c) mean brain signal within the lateral ventricles, and (d) the mean signal within a deep white matter ROI. Inclusion of the first temporal derivatives of these regressors within the linear model accounted for the time-shifted versions of spurious variance. Global signal regression was included, which uses a general linear model to regress out the average signal across all voxels in the brain, including physiological noise (e.g. cardiac and respiratory), movement-related artifact, and nonspecific signals. The time course of the average blood-oxygen-level-dependent (BOLD) signal within each spherical ROI was compared with the BOLD signal time course of other brain voxels to identify regions with positive and negative correlations. Pearson correlation coefficients were converted to normally distributed Z-scores using the Fisher transformation. To identify the functional network that best represents the individual rs-fcMRI maps we employed a weighted principal component analysis in MATLAB. Images were created using 3D Slicer, ImageMagick, Surf Ice, and GIMP.

To identify common cognitive impairments that co-occur with time disorientation, the distribution of scores for neuropsychological tests available for at least 75% of the time disoriented patients was compared to the distribution of scores in the brain-damaged comparison group. The scores were not normally distributed, so we used Quade’s Ranked ANCOVA (RANCOVA) [37] to compare tests with continuous scores and Pearson’s Chi Square Test for categorical scores. Several of the cognitive tests used in our analyses were subtests from the WAIS, the most widely used measure of adult intelligence [38]. If a participant was administered multiple versions of the WAIS, the most recent version was used [39].

We performed a post hoc analysis of lesion location for time disoriented participants with relatively intact cognitive performance, which we defined as being in the normal range (not more than 1 standard deviation below the mean) on >75% of the neuropsychological tests included in this study. Lesion location was evaluated in this subgroup analysis using proportional subtraction as the sample size was underpowered for a LESYMAP analysis. Participants were divided into impaired (BTOT score of -3 or worse) and unimpaired (BTOT score of 0) groups. The proportional metric (PM) was calculated using FSL [40] using previously described methods [41]. The proportion of time disoriented patients with a lesion relative to all time disoriented patients was calculated on a voxel-wise basis. The same proportional map was created for time oriented patients and a difference map of the impaired and unimpaired proportional maps was generated such that voxels were weighted from -1 to 1, with higher scores indicating a stronger relationship between damage at that voxel and impairment on the BTOT.

### Data availability

Anonymized data not published within this article will be made available by request from any qualified investigator.

## Results

A total of 39 individuals with time disorientation were identified. Relative to the brain-damaged comparison group, on average those with time disorientation were older (p < .001) had larger lesions (p < .001), a higher rate of bilateral lesions (48.82% versus 19.18%), and a higher rate of encephalitis (12.82% versus 1.37%, p<.001). In the impaired group, 5 patients had encephalitis, 9 patients had a stroke of the posterior artery distribution (3 left hemisphere, 3 right hemisphere, and 3 bilateral). 7 patients had lesions in the frontal lobe, mostly involving the ventromedial prefrontal cortex, 2 lesions involved the thalamus and intersected with the mediodorsal nucleus of the thalamus, and 15 lesions involved posteromedial cortices. Lesions of the 39 individuals with time disorientation are shown in *Figure 1a*. This distribution can be compared to the lesion masks of all 550 participants in this study, which spanned virtually the entire cortex. Of the 550 participants, a maximum overlap of 51 lesions was observed in the right frontal white matter (*Figure 1b*). The overlap map of the 75 patients with BTOT scores of -1 and -2 that were not included in the main analysis showed a similar distribution to the brain-damaged comparison group (*Figure 1c*).

**Figures 1a-c.**
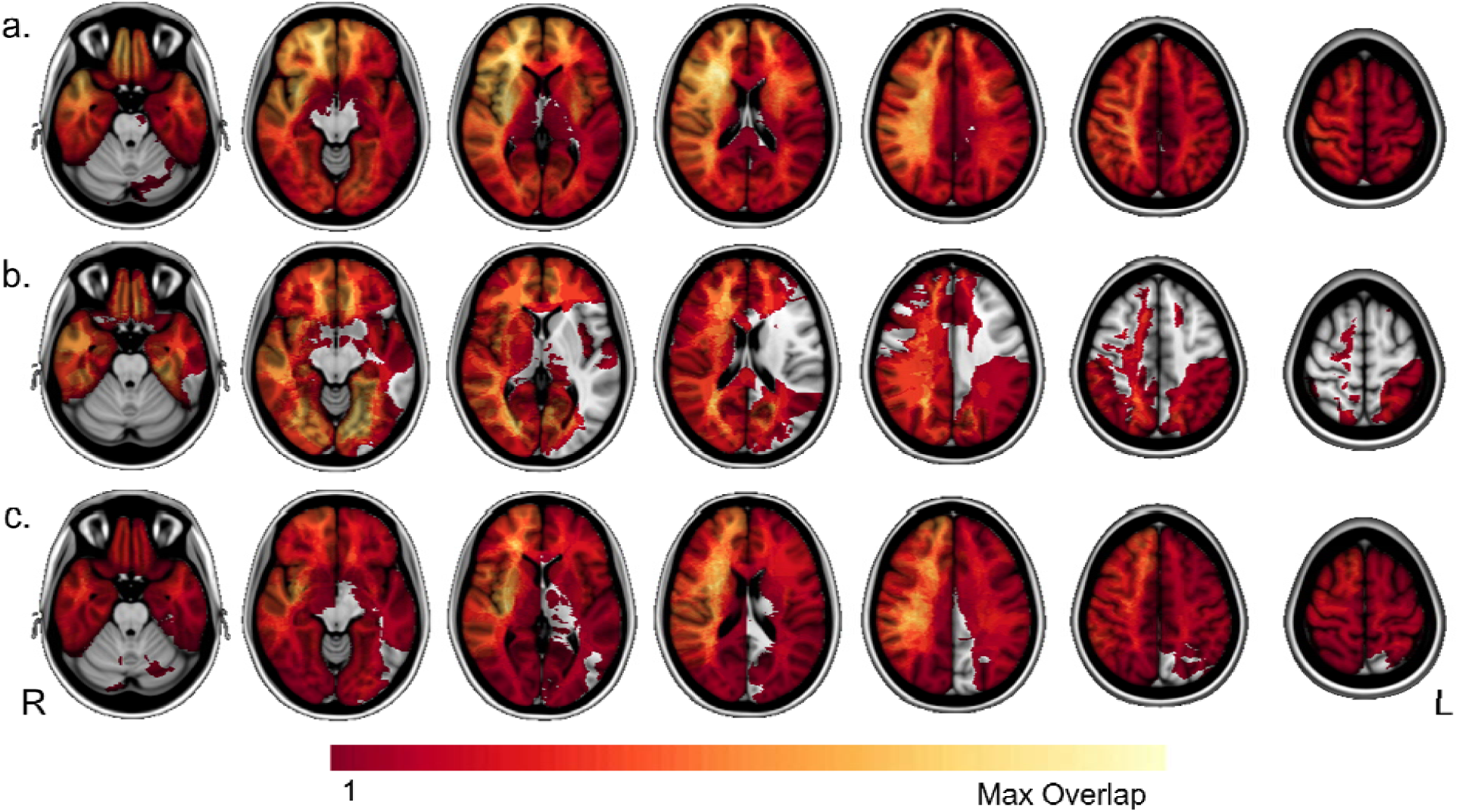
Lesion masks of all 550 participants in this study were overlapped to show that we have lesion coverage across a majority of the brain to assess brain-behavior relationships. The maximum number of overlapping lesions (51) was located in right frontal white matter (*1a*). The lesion masks of the 39 time disoriented participants were also overlapped (*1b*); the region of peak overlap in the left parahippocampal gyrus contained 9 lesions. Of the patients tested, 75 individuals earned a score of one or two on the BTOT. The distribution of these patients’ lesions is similar to the unimpaired group. The voxel of peak lesion overlap (19 lesions) is also in right frontal white matter (*1c*).

### Lesion-Symptom Mapping Results

The lesion-symptom mapping of all 550 participants was significant at r=.22, p<.001. The peak regional findings occurred at the medial temporal cortex (MNI -36 -35 -21), the precuneus (−20 -67 25), and white matter between them (−22 -74 15), in addition to visual areas V2 (−13 -73 -3) and V4 (−25 -79 -16) (*Figure 2a*). The analysis of unilateral lesions (n=433) was only significant when all lesions were represented on the same hemisphere (r=.21, p<.001), with localization in the precuneus (−20 -67 25), parahippocampal gyrus (−42 -31 -25), and white matter between these regions (−18 -78 15) (*Figure 2b*). The main difference between the lesion-symptom maps of all subjects versus those with unilateral lesions was retention of the medial temporal lobe and precuneus findings with unilateral lesions with less occipital lobe findings upon removing individuals with bilateral lesions.

**Figures 2a-c.**
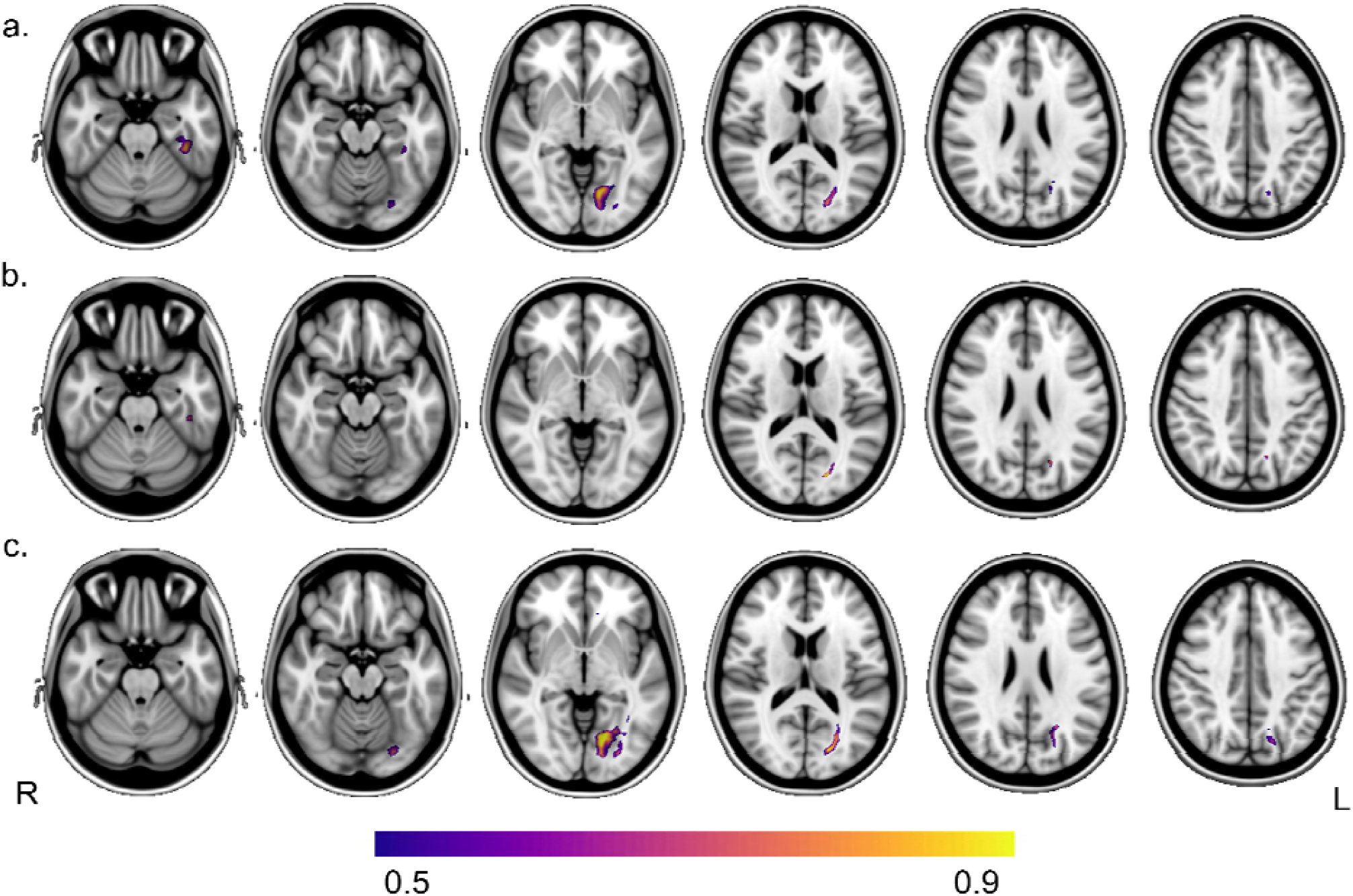
The lesion-symptom mapping analysis was first conducted with the entire sample (*2a*) and showed regions of peak intensity in the precuneus, medial temporal lobe, and visual areas V2 and V4. To address the impact of focal unilateral lesions more specifically, a lesion-symptom mapping analysis including only unilateral lesions showed regions of peak intensity in the precuneus, parahippocampal gyrus, and other areas of the MTL (*2b*). Results from both analyses reveal a white matter region possibly connecting the precuneus and MTL, is important for time orientation too. After removing 12 participants with lesions caused by encephalitis (n=538), peak regions in the occipital lobe, precuneus, and white matter tract previously identified were evident (*2c*).

Lesion network maps were derived from the peak regional findings from the lesion-symptom maps. The associated networks were used as input to a principal component analysis to identify networks that capture variance across these regional findings. The two main components resembled visual and default mode networks, respectively (*Figures 3a&b*). The precuneus site, in particular, appeared to drive the 2^nd^ principal component, which showed functional connections to the occipital lobe and MTL (*Figure 3c*).

**Figures 3a-c.**
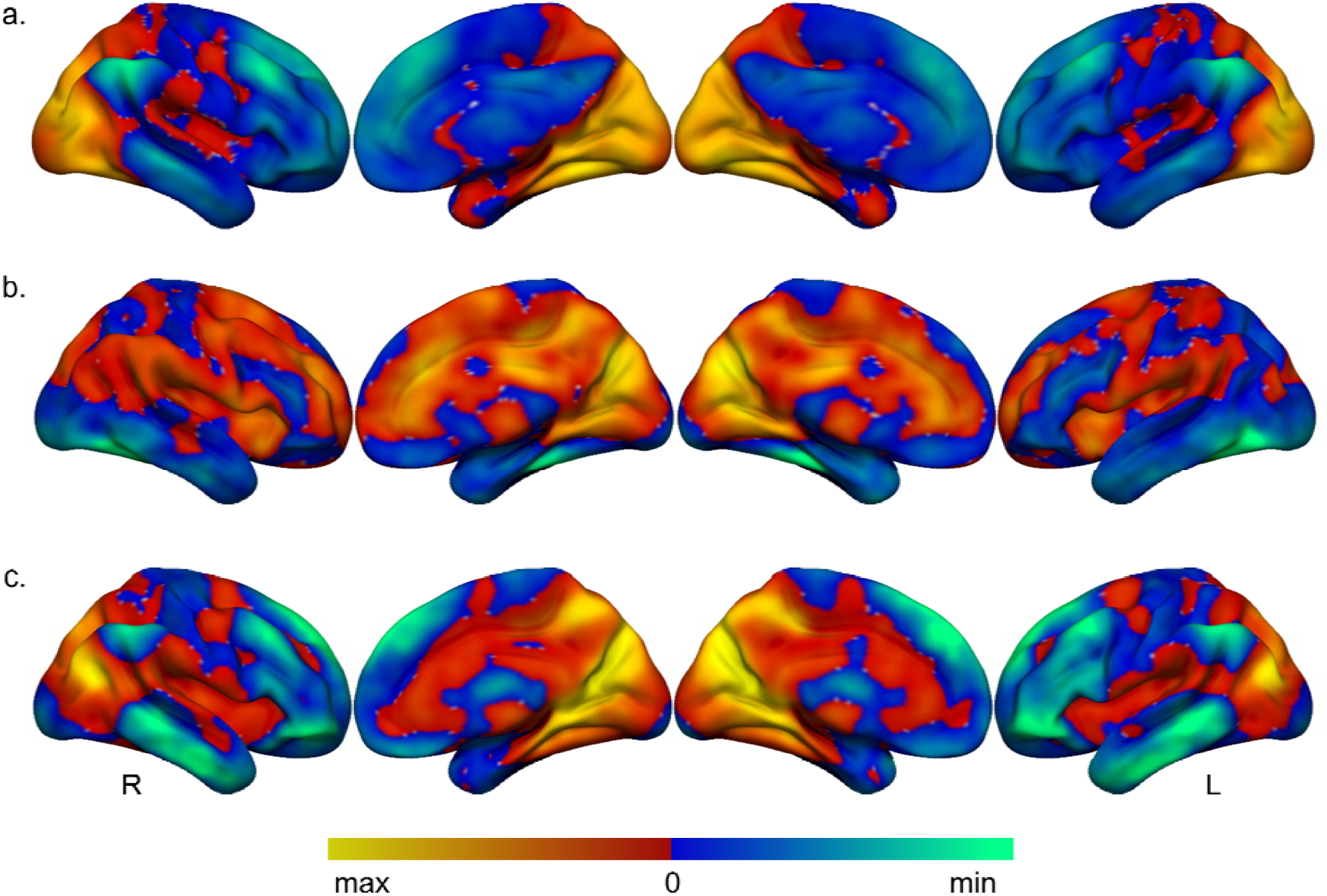
A principal component analysis of the lesion network maps seeded by the four peak regions from the LESYMAP reveals regions within the visual network (min=-8, max=10) (*3a*) and default mode network (min=-6, max=6) (*3b*) explain most of the variance in the data. A seed ROI placed over the peak voxel in the precuneus seeded a lesion-derived network map (min=-6, max=12) (*3c*) which showed functional connectivity between precuneus, MTL, and occipital regions.

To characterize cognitive impairments that co-occur with time disorientation, we compared the distribution of scores between the impaired (n=39) and brain-damaged comparison (n=511) groups across a variety of neuropsychological tests (*Table 3*). Age and lesion volume were included as covariates in our RANCOVA given significant between-group differences. After correcting for multiple comparisons, the Bonferroni corrected alpha was .002. The group with time disorientation performed significantly worse on memory tests, including the Rey Auditory-Verbal Learning Test (verbal memory), Rey-Osterrieth Complex Figure Test (visuoconstruction and visuospatial memory), as well as naming (Boston Naming Test), and processing speed (Coding subtest of WAIS).

**Table 3.**
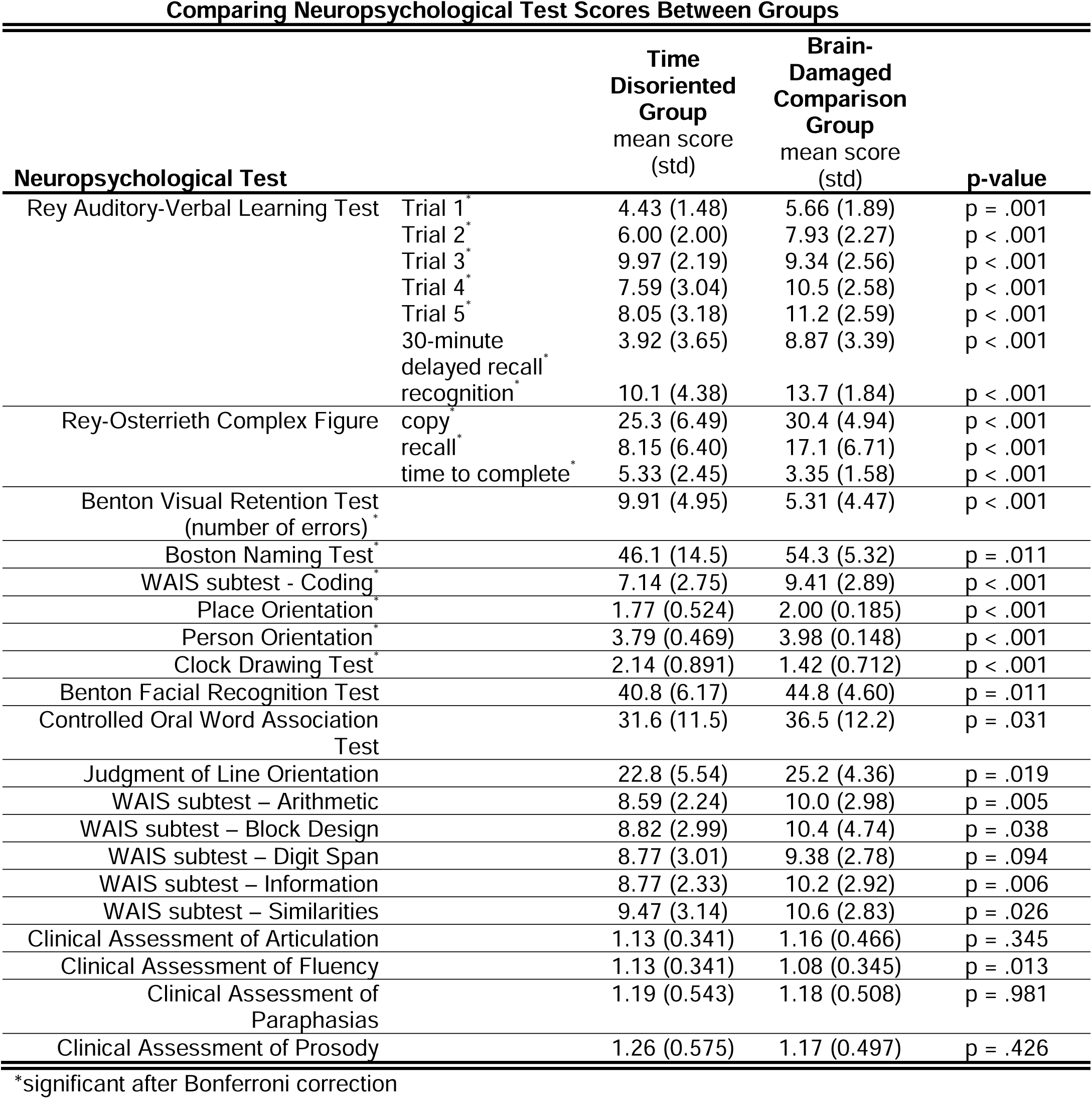
We performed Quade’s Ranked ANCOVA using age and lesion volume as covariates to compare the distribution of neuropsychological test scores between the time oriented and time disoriented groups.

Finally, based on the findings above of significant differences in cognitive performance in the individuals with impaired time orientation, a post-hoc analysis was performed. We aimed to address an alternative explanation that our neuroanatomical findings of the lesion-symptom maps may reflect memory impairment or global cognitive impairment rather than our primary function of interest, time disorientation. We identified 11 time disoriented patients with relatively preserved cognition (defined as impaired on fewer than 25% of the neuropsychological tests in *Table 3*). A proportional subtraction analysis was performed on this cohort with a peak regional finding at the left precuneus (*Figure 4a*).

**Figures 4a&b.**
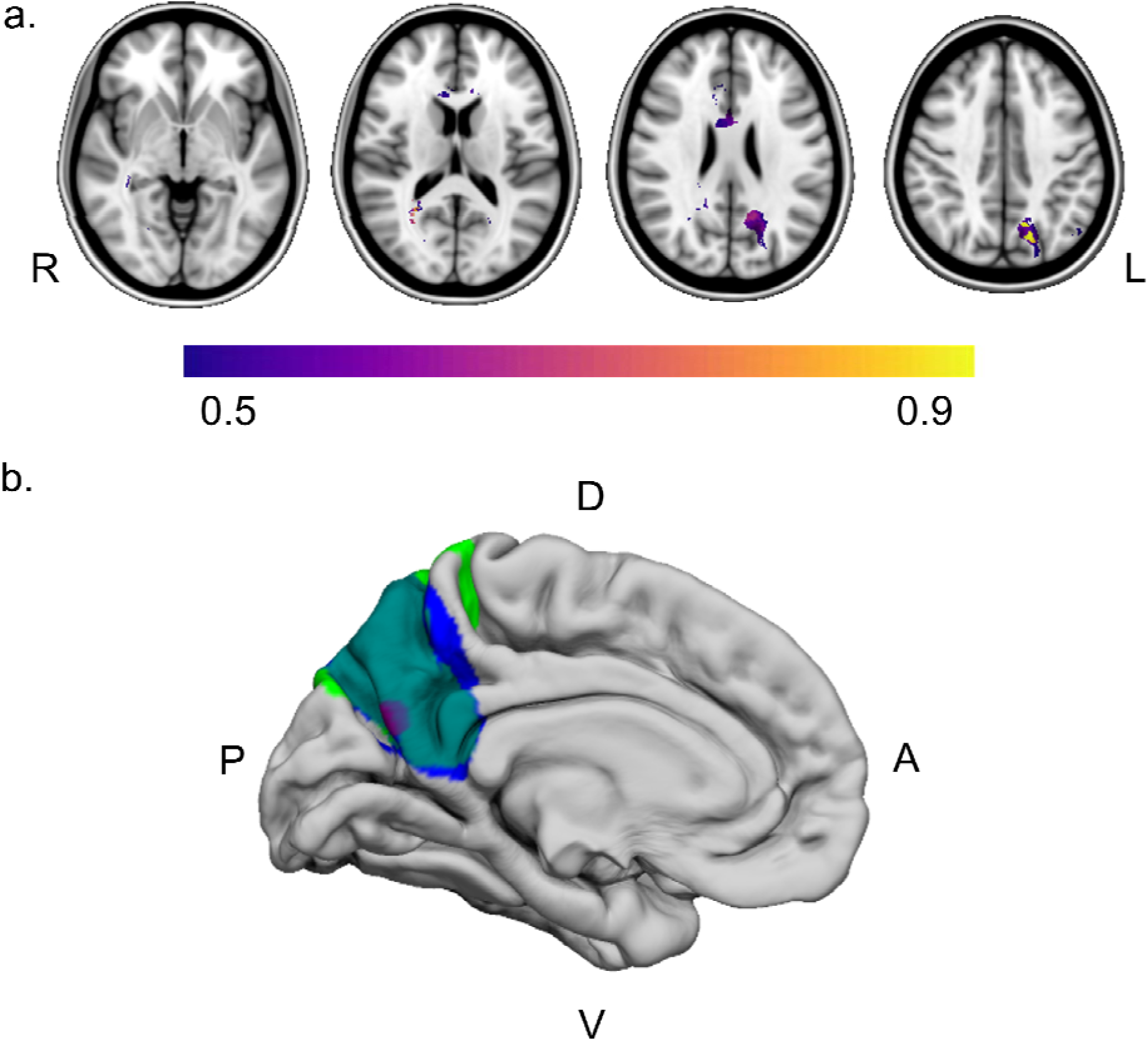
A proportional subtraction analysis of 11 time disorientated patients with relatively unimpaired cognitive performance in other domains was conducted (PM=0.96 at location -13, -61, 37 in the precuneus) (*4a*). The lesion location of two individuals with tumor resection surgeries are shown (in blue and green, with overlap of the two lesions in turquoise) (*4b*). Both individuals had lesions that overlapped with our lesion-symptom mapping results in the precuneus and spontaneously reported major alterations in their experience of time following the lesion. The lesion-symptom mapping result is just deep to the cortical surface so a purple marker is shown to denote the location.

### Subjective Experience of Time Following Precuneus Lesions

In light of our lesion-symptom mapping findings in the precuneus (*Figure 2*) we performed a post-hoc review of medical records of 16 individuals following tumor removal that resulted in a focal lesion of the precuneus. This cohort was described in detail previously [42]. Time orientation was not systematically assessed in this retrospectively analyzed cohort, yet two individuals with left precuneus lesions that overlap with the lesion-symptom map findings (*Figure 4b*) spontaneously reported a notable change in their experience of time. One patient reported that “time did not run” and experienced subjective time dilation, experiencing minutes as hours. Days felt long to him and the time on his watch was systematically earlier than he expected. The second patient also spontaneously reported that “time did not run” and noted difficulties conceptualizing time, with his clinical notes documenting trouble remembering the current date. While these two reports are anecdotal they are aligned with the lesion-symptom map in supporting a role of the precuneus in influencing time perception and time orientation.

## Discussion

The main findings of our analysis are that damage to the MTL, precuneus, and regions of the ventromedial occipital lobe is associated with time disorientation. Other notable findings of interest include the observation of multiple risk factors for time disorientation, including: bilateral lesions, older age, lesions from encephalitis (as has been reported previously {Greenwood, 1983 #83). Somewhat unexpectedly, lesions involving the posterior cerebral artery distribution were more associated with time disorientation relative to prefrontal cortex lesions (n=9 versus n=7).

The precuneus result is of particular interest because it is present in analyses limited to subjects with unilateral lesions and it is the peak regional finding in an analysis of subjects with relatively preserved overall cognitive performance. As such, it may be more specific to time orientation relative to other regional findings that occurred with bilateral lesions or co-occurred with cognitive deficits in other domains. Additional support for the possibility of precuneus involvement in time orientation came from a post-hoc analysis that identified two patients from an independent cohort with focal lesions involving this left precuneus region that spontaneously reported major distortions in their perception of time in the immediate days following the lesion onset, both reporting a subjective prolongation of their subjective experience relative to clock time. The passing of minutes felt more like hours. The functional connectivity network associated with this precuneus site is most closely aligned with the default mode network, which includes medial temporal regions that were also implicated in time disorientation.

The precuneus has been implicated in time-related cognition from functional imaging studies, but this association with time disorientation has not been observed in lesions studies to our knowledge. Focal lesions to the precuneus are extremely uncommon [43]. The precuneus is a highly connected region associated with complex information integration and has a high resting metabolic rate [44]. It is known to be involved in self-referential processes broadly [44], including autobiographical memory [45]. In terms of time-related processing, an fMRI study show activation in the precuneus in longer-scale temporal ordering tasks [46] and time orientation in healthy humans [17]. Differences in precuneus activity were observed in relation to a time reproduction task in medicated and non-medicated patients with Parkinson’s disease [47].

Peaks in the parahippocampal gyrus and surrounding medial temporal cortex are in line with prior research suggesting that damage to and dysfunction of the MTL is associated with time disorientation across various etiologies [7, 11, 14].

We also observed regional findings in the ventromedial occipital lobe and extending anteriorly along the temporo-occipital junction. Our working interpretation of these findings is they result from lesions that disrupt communication between the other regional findings in the precuneus and MTL, yet we cannot rule out the possibility that this region is also important in time orientation independent of these other sites. There are proposals that vision and time-related processing may be interrelated, and our results are consistent with this possibility [44, 48, 49].

Our study has limitations. First, while our overall sample size was large there were only 39 individuals with time disorientation. A larger sample size with chronic time disorientation will be useful in further refining the underlying anatomy. Second, chronic time disorientation can wax and wane over time. A patient may be more or less disoriented at the time of testing and since test scores most contemporaneous with the scan date were utilized, scores could vary. But, orientation to time typically does not oscillate between disoriented and oriented, just in the degree of impairment [50]. Next, different time scales related to one’s mental timeline may be represented differently in the brain. Long-term memory required for remembering the year may involve different brain regions compared to those important for shorter time scales like knowing the time on a clock. A future analysis of the neuroanatomical correlates of time disorientation and varying time-scales could address this limitation. Finally, we did not have good lesion coverage in some brain areas that may be relevant for time orientation, such as the thalamus and cerebellum. Prior research indicates that damage to the mediodorsal nucleus of the thalamus is associated with time disorientation, also known as thalamic chronotaraxis [14, 15] and the cerebellum may also have a role [51].

This study begins to formulate a neuroanatomical network important for time orientation. In doing so it aligns with prior work implicating lesions of the medial temporal lobe, and we extend this work in also implicating the precuneus and medial occipito-temporal regions. With regard to how the human brain represents time there are many outstanding questions that await future studies, including the neuroanatomy of other types of timing such as sub- and supra-second interval timing. We are optimistic a better understanding of these processes will continue to inform how the brain creates a temporal context for events and how these processes are utilized to support different domains of cognition.

## SUPPLEMENTAL FIGURE 1

*Supplemental Figure 1a-c*. Peak regions from the LESYMAP analysis in the parahippocampal area (*a*) and occipital lobe (*b&c*) seeded lesion network mapping analyses. The results overlap most with the default mode network and the visual network.

## Notes

### Competing Interest Statement

The authors have declared no competing interest.

