## Supplementary figures and images for "Lesion Localization of Time Disorientation in Patients With Focal Brain Damage"

### Supplemental Figure 1

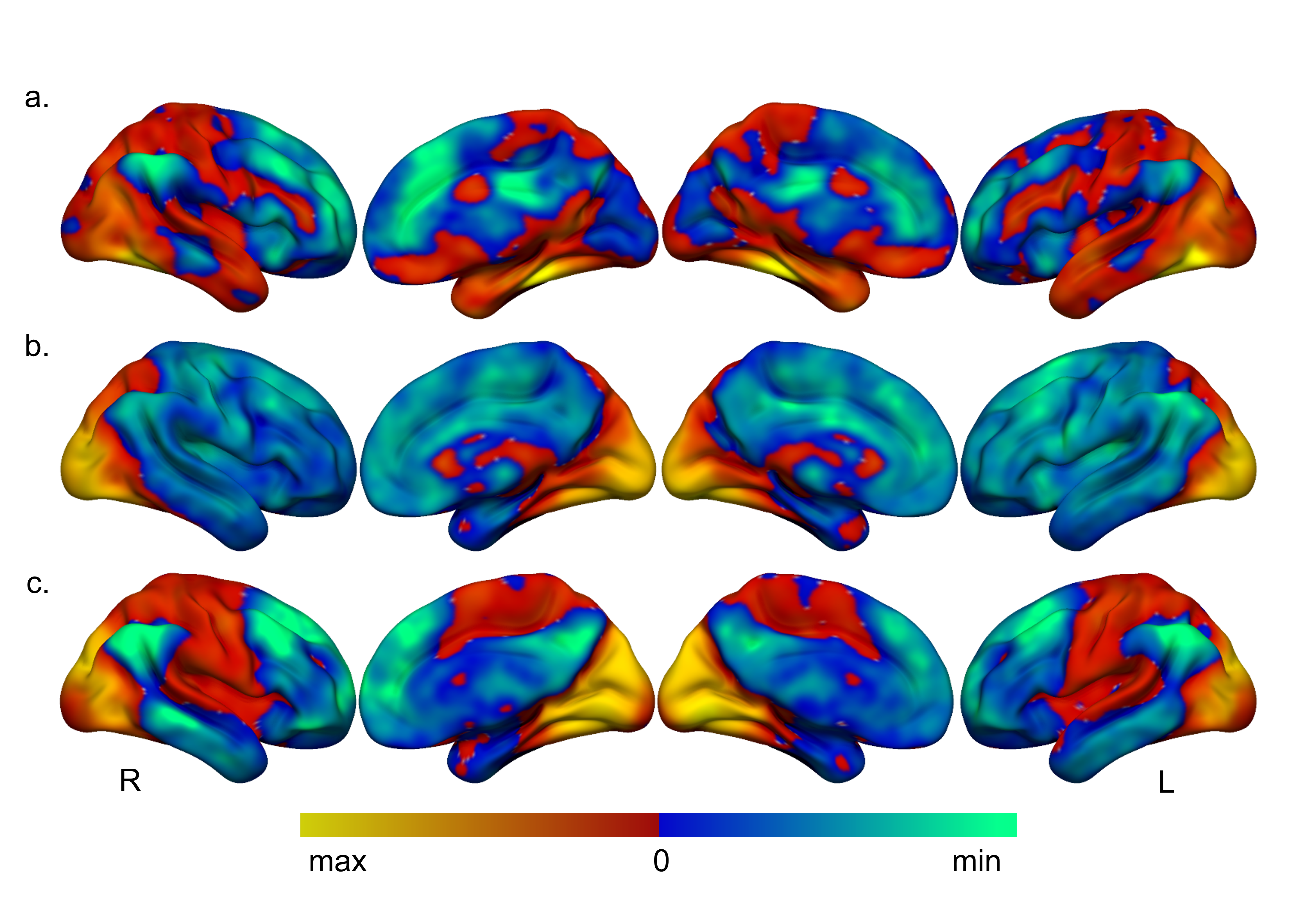
